# Structure-based design and characterization of Parkin activating mutations

**DOI:** 10.1101/2022.02.22.481412

**Authors:** Michael U. Stevens, Nathalie Croteau, Mohamed A. Eldeeb, Zhi Wei Zeng, Rachel Toth, Thomas M. Durcan, Wolfdieter Springer, Edward A. Fon, Miratul M. K. Muqit, Jean-François Trempe

**Author notes:** Equal contributions (Michael U. Stevens, Nathalie Croteau, Mohamed A. Eldeeb).

## Abstract

Human autosomal recessive mutations in the Parkin gene are causal for Parkinson’s disease (PD). Parkin encodes a ubiquitin E3 ligase that functions together with the PD associated kinase, PINK1, in a mitochondrial quality control pathway. Structural studies reveal that Parkin exists in an inactive conformation mediated by multiple autoinhibitory domain interfaces. Here we have performed comprehensive mutational analysis of both human and rat Parkin to unbiasedly determine Parkin activating mutations across all major autoinhibitory interfaces. Out of 31 mutations tested, we identify 11 activating mutations clustered near the RING0:RING2 or REP:RING1 interfaces, which reduce the thermal stability of Parkin. Of these, we demonstrate that three mutations, V393D, A401D, and W403A located at the REP:RING1 interface were able to completely rescue a Parkin S65A mutant, defective in mitophagy, in cell-based studies. Overall our data extends previous analysis of Parkin activation mutants and suggests that small molecules that mimic REP:RING1 destabilisation offer therapeutic potential for PD patients harbouring select Parkin mutations.

**Summary blurb:** Parkin, an E3 ubiquitin ligase involved in Parkinson’s disease, is inactive in the basal state and is activated by PINK1 to mediate mitophagy. Here we characterized 31 mutations and discovered three that activate Parkin and rescue loss of PINK1 phosphorylation.

## Introduction

Mutations in the *PRKN* (Kitada et al 1998) and *PINK1* (Valente et al 2004) genes account for the majority of recessive early-onset Parkinson’s Disease (PD) cases (Martin et al 2011). The *PRKN* gene encodes Parkin, an E3 ubiquitin (Ub) ligase that mediate a mitochondrial quality control pathway that is also dependent on the PINK1 kinase activity (Bayne & Trempe 2019). Evidence in support of this hypothesis initially came from genetic models in *Drosophila*, where ablation of either genes causes muscle degeneration and male sterility as a result of mitochondrial defects (Clark et al 2006, Greene et al 2003, Park et al 2006). While *PRKN* knockout (KO) mice do not display neuron loss (Goldberg et al 2003), they exhibit deficits in striatal neuron excitability, as well as perturbation in mitochondrial proteins and respiration (Palacino et al 2004, Periquet et al 2005). Genetic crosses of *PRKN* KO with the *mutator* mouse, which has a defective mitochondrial DNA polymerase, show nigrostriatal degeneration and inflammation, consistent with Parkin having a role in protecting Dopaminergic (DA) neurons from mitochondrial damage (Pickrell et al 2015, Sliter et al 2018). Likewise, *PINK1* KO mice also do not display neurodegeneration, but similar *to PRKN KO* exhibit marked mitochondrial defects (Gautier et al 2008). Furthermore, *PINK1* KO mice subjected to bacterial infections undergo loss of DA axonal varicosities, as a result of mitochondrial antigen presentation and activation of cytotoxic T cells in the brain (Matheoud et al 2019). Thus, loss of the Parkin or PINK1 proteins results in mitochondrial dysfunction, which can ultimately lead to neurodegeneration. It has also been reported that the levels of soluble Parkin decrease with age and PD patients show elevated levels of insoluble Parkin, which would be associated with reduced biochemical activity (Pawlyk et al 2003, Tokarew et al 2021, Wang et al 2005). Therefore, enhancing Parkin’s intrinsic activity may have beneficial therapeutic implications

The mechanism of action of Parkin and PINK1 has been elucidated in the last decade. In cultured cells, the two proteins mediate selective mitochondrial autophagy (mitophagy) following chemical-induced mitochondrial depolarization; accumulation of misfolded mitochondrial proteins; or following exposure to complex I inhibitors (Geisler et al 2010, Jin & Youle 2013, Matsuda et al 2010, Narendra et al 2008, Narendra et al 2010). PINK1 is a kinase stabilised on damaged mitochondria, where it phosphorylates ubiquitin at Serine65 (Kane et al 2014, Kazlauskaite et al 2014, Koyano et al 2014). The resulting phospho^Ser65^-ubiquitin (pUb) acts as a receptor for Parkin to be recruited to sites of damaged mitochondria, thus enabling PINK1 to phosphorylate Parkin, which in turn dramatically increases its E3 ubiquitin ligase activity (Kondapalli et al 2012). Active Parkin ubiquitinates substrates on the outer mitochondrial membrane (OMM) such as Mitofusin, Miro, VDAC, etc, which are modified at different rates and extent (Antico et al 2021, Chan et al 2011, Ordureau et al 2018, Tanaka et al 2010, Vranas et al 2022). The accumulation of the ubiquitinated proteins is rapid due to a feedforward mechanism and has mainly been linked to mitophagy (Ordureau et al 2014). Beyond mitophagy, Parkin/PINK1 can also mediate mitochondrial arrest in neurons (Wang et al 2011), spatially-restricted mitophagy (Yang & Yang 2013), and mitochondria-derived vesicle (MDV) formation (McLelland et al 2014). Parkin/PINK1 most likely mediate stress-evoked degradation of a subset of mitochondrial proteins, as evidenced by turnover measurements in *Drosophila* (Vincow et al 2013), and which is distinct from the regulation of basal mitophagy that is independent of PINK1 in mice (McWilliams et al 2018b). Furthermore, pUb can be detected in human and rodent brains (Fiesel et al 2015), is absent in PINK1-null mammals and reduced in Parkin-null or Parkin^S65A^ mutant knock-in mice, and is increased in the mutator mouse and sporadic PD patients (Hou et al 2018, McWilliams et al 2018a, Pickrell et al 2015, Watzlawik et al 2021). These observations strongly support PINK1 being the primary activator of Parkin in vivo, with pUb serving as a marker for mitochondrial damage.

Structural studies have unveiled fundamental details regarding Parkin activation at the molecular level. Parkin belongs to the RING-in-Between-RING (RBR) family of E3 Ub ligases, which transfers ubiquitin to a substrate from an E2 ubiquitin-conjugating enzyme via formation of a thioester intermediate on a reactive cysteine (Parkin Cys431) in the RING2 catalytic domain (Wenzel et al 2011). Structural and biochemical analysis of Parkin showed that the protein is auto-inhibited in the basal, unstimulated state (Chaugule et al 2011, Riley et al 2013, Spratt et al 2013, Trempe et al 2013, Wauer & Komander 2013). The ubiquitin-like (Ubl) domain and Repressor Element of Parkin (REP) both occlude the E2-binding site on RING1, and Cys431 is sequestered at the interface of RING0 and too far from the E2∼Ub conjugate for efficient transfer. Upon binding to pUb, the Ubl domain is released from RING1 and can be freely phosphorylated by PINK1 at Ser65 (Kazlauskaite et al 2015, Kumar et al 2015, Sauvé et al 2015, Wauer et al 2015). The phospho-Ubl domain then binds to RING0 and competes with the RING2 domain, which dissociates concomitantly with the REP, thus allowing E2∼Ub binding (Condos et al 2018, Gladkova et al 2018, Sauvé et al 2018). While there is no crystal structure of the resulting Parkin thioester transfer complex, how the RING2 domain will interact with E2∼Ub, can be inferred from the structure of the RBR ligase HOIP bound to UbcH5B∼Ub (Lechtenberg et al 2016). Once Ub is charged on Cys431, it is transferred to a substrate lysine in a reaction catalysed by His433 (Vranas et al 2022). The broad range of Parkin substrates suggest that is unlikely to recognize a specific substrate motif, and substrate specificity is rather conferred by proximity at sites of activation.

In the light of these *in vitro* studies, it appears that interdomain interactions maintain Parkin in an auto-inhibited state. Indeed, mutagenesis of residues located at inhibitory interfaces, such as Phe146 in RING0 or Trp403 in the REP, result in a dramatic increase in the ligase activity of Parkin and mitochondrial recruitment (Trempe et al 2013), and can rescue S65A or *Δ*Ubl Parkin in ubiquitination and mitophagy assays (Sauvé et al 2015, Tang et al 2017). Synthetic F146A and W403A mutations also rescue seven PD-associated Parkin missense mutations that disrupt Parkin activity through various mechanisms (Yi et al 2019). On the other hand, the N273K mutation in RING1, which repels the Ubl domain, accelerates mitochondrial recruitment but does not rescue the S65A mutation (Tang et al 2017). Since activating mutations can result in the constitutive activation of Parkin in the cytosol, it is conceivable that strongly activating mutations may induce rapid Parkin turnover via auto-ubiquitination, which would reduce the steady-state levels of the protein in cells and thus reduce its actual activity towards mitochondrial substrates. Another factor to consider is the effect of the mutation on the solubility of Parkin; for instance, we have found that mutations at the RING0:RING2 interface reduce the solubility of the protein *in vitro* (Tang et al 2017). Finally, only a small subset of interfaces and mutants has so far been tested, each of which may tune Parkin’s activity and stability to various degrees. Thus, a systematic and comprehensive assessment of activating mutations is required.

Here, we have designed and introduced 31 synthetic mutations in mammalian Parkin as candidates for activation, and have characterized their effects on recombinant protein expression, thermal stability, and ubiquitination activity in vitro. We identified 11 mutations that activate both rat and human Parkin. We find that all activating mutations cluster near the RING0:RING2 or REP:RING1 interfaces and reduce the thermal stability of Parkin. These mutations were introduced in human cells and tested for their ability to enhance depolarization-induced mitophagy using a fluorescent-reporter assay. From this group of 11, we identified two novel mutations (V393D and A401D) that robustly enhance mitophagy and rescue the defective S65A mutation, in addition to the W403A mutation previously reported. These mutations provide a molecular basis for future studies in neuronal cells and map to a “hot-spot” of activation in Parkin that will serve as a scaffold for the design of small-molecule activators.

## Results

### Survey and design of activating mutations in Parkin

Different groups have reported mutations that activate Parkin in vitro (Table 1). These include mutations at the RING0:RING2 and REP:RING1 interfaces (F146A, W403A, F463Y, F463A) (Riley et al 2013, Trempe et al 2013, Wauer & Komander 2013), as well as residues within the RING0:RING1 interface that remodel upon phosphorylation (H227N, E300A/Q) (Kumar et al 2015). Mutations in RING1 residues that bind to the Ubl (N273K, L266K/R) also activate Parkin (Kumar et al 2015, Sauvé et al 2015, Wauer et al 2015). While mutations in the Ubl at the corresponding interface also activate Parkin in vitro (I44A, H68A, etc.) (Kumar et al 2015), these mutants are likely to interfere with PINK1 phosphorylation (Rasool et al 2018, Wauer et al 2015), and were avoided. The PD mutation R33Q has also been shown to activate Parkin (Chaugule et al 2011).

**Table 1.** Proposed activating mutations in Parkin (conserved in rat and human Parkin)

| **Mutation** | **Domain** | **Reference(s)** |
| --- | --- | --- |
| R33Q | UBL | (Chaugule et al 2011) |
| Y143D | RING0 | This paper |
| Y143E | RING0 | This paper |
| F146A | RING0 | (Ordureau et al 2014, Tang et al 2017, Trempe et al 2013, Yi et al 2019) |
| W183Y | RING0 | This paper |
| F208Y | RING0 | This paper |
| H227N (human only) | RING1 | (Kumar et al 2015) |
| H/N227A (human/rat) | RING1 | This paper |
| L266K | RING1 | (Sauvé et al 2015) |
| L266R | RING1 | This paper |
| N273K | RING1 | (Sauvé et al 2015, Tang et al 2017) |
| E300A | RING1 | (Kumar et al 2015) |
| E300Q | RING1 | (Kumar et al 2015) |
| E370A | IBR | This paper |
| V393D | REP | This paper |
| V393K | REP | This paper |
| A397D | REP | This paper |
| A397K | REP | This paper |
| A398T | REP | (Periquet et al 2003) |
| A398D | REP | This paper |
| A398K | REP | This paper |
| A401D | REP | This paper |
| A401K | REP | This paper |
| W403A | REP | (Ordureau et al 2014, Sauvé et al 2015, Tang et al 2017, Trempe et al 2013, Yi et al 2019) |
| W403F | REP | This paper |
| W462A | RING2 | This paper |
| W462F | RING2 | This paper |
| W462Y | RING2 | This paper |
| F463Y | RING2 | (Riley et al 2013) |
| D464A | RING2 | This paper |
| D464K | RING2 | This paper |

In addition to the mutations reported above, we also sought to design new mutations that may increase Parkin’s activity based on structures of inactive and active mammalian Parkin (Figure 1). Since human His227 is an Asn in rat and mouse Parkin, this residue was mutated to Ala as well (Table 1). This residue is located in proximity to Trp183 at the RING0:RING1 interface, and thus we also introduced the W183Y mutation. Given that the REP W403A mutant displays some of the most desirable characteristics (stable, soluble, enhances Parkin recruitment and mitophagy), we introduced multiple additional REP mutations (Figure 1). In particular, we mutated small aliphatic residues in the REP helix to charged residues in order to disrupt their interactions with the RING1 domain; namely residues Val393, Ala397, Ala398 and Ala401 (Figure 1). Intriguingly, A398T was discovered in a single PD patient as a heterozygous mutation (Periquet et al 2003), but it is unclear whether it is pathogenic. It was also reported to activate Parkin in vitro (Wauer & Komander 2013), and we therefore included this mutation in our panel. In addition, we mutated Trp403 to a Phe to potentially achieve a lower degree of activation compared to W403A. We noted that Trp462 in RING2 is also mediating an interaction with Phe208 in RING0, and therefore mutated these aromatic residues to Tyr to maintain aromaticity but perturb the interaction (Figure 1). Finally, two groups reported that Abl phosphorylates Parkin on Tyr143, which was reported to inactivate the enzyme (Imam et al 2011, Ko et al 2010). However, inspection of the rat apo Parkin structure suggests that introducing a negative charge at this position may induce an electrostatic repulsion with the C-terminus of RING2 and thus may result in the activation of Parkin (Figure 1). We thus introduced the Y143D and Y143E mutations.

**Figure 1.**
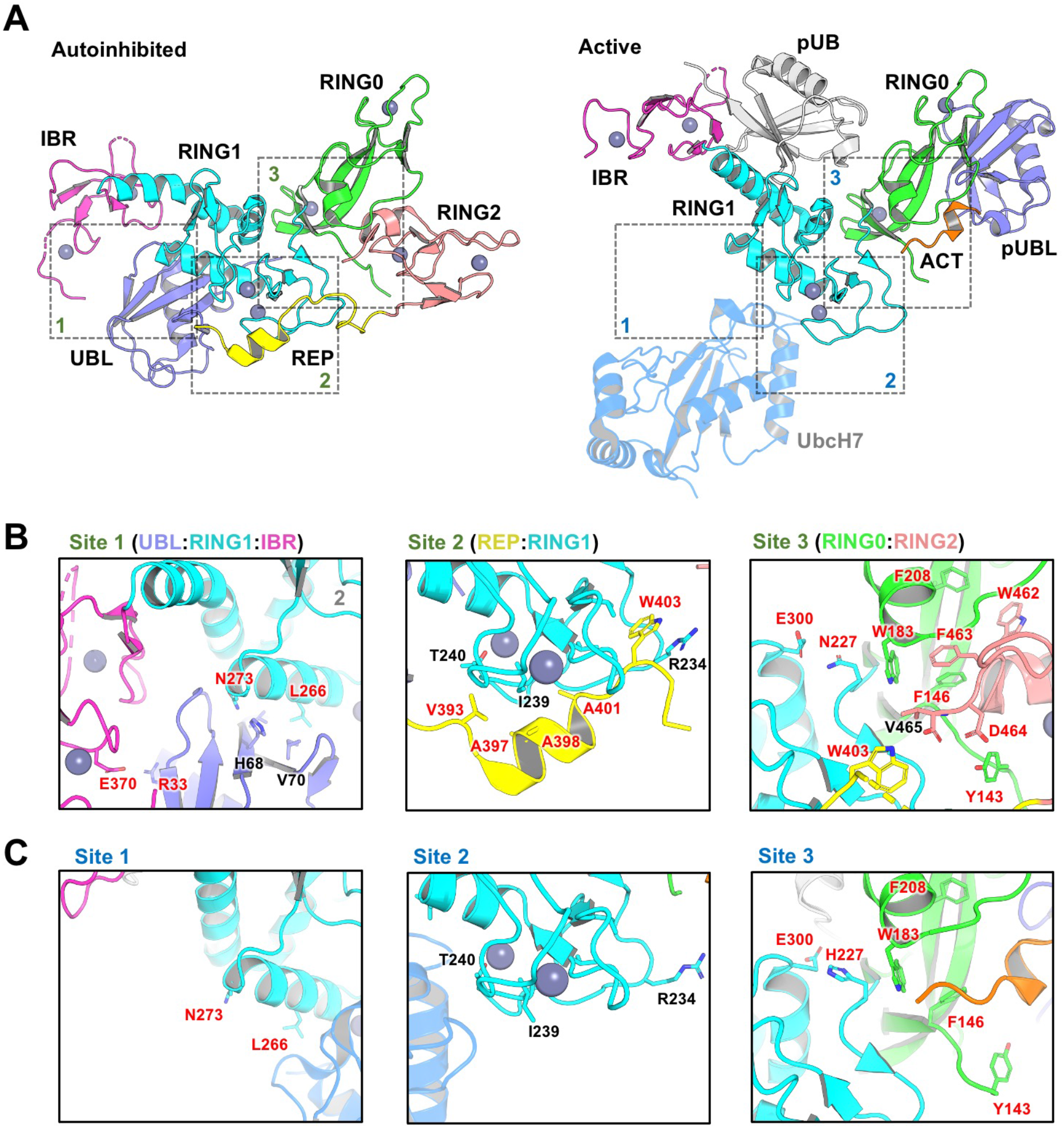
Design of activating mutations in Parkin. **(A)** Cartoon representations of autoinhibited apo rat Parkin (left; PDB ID: 4ZYN) and active human phospho-Parkin bound to phospho-Ub (right; PDB ID: 6GLC). The structure of autoinhibited apo human Parkin (PDB ID 5C1Z, not shown) is nearly identical to the rat ortholog but lacks Y143. The position of UbcH7 is based on the superposition of activated fly phospho-Parkin bound to pUb and UbcH7 (PDB ID: 6DJW, not shown). The different domains of Parkin are shaded and labelled, and dashed boxes indicate interfaces between these domains. **(B)** Residues within the binding interfaces of auto-inhibited rat Parkin are highlighted, with residues mutated for this work labeled in red. **(C)** Residues within the same binding interfaces as in B, but for the activated human phospho-Parkin:pUb complex.

All mutations mentioned above were introduced in both human and rat Parkin, to identify mutations that could later be studied in both rodent models and human cells. Furthermore, the experiments with rat and human Parkin were performed in two different labs using different plasmids for bacterial expression, to ensure robust identification of mutations that activate Parkin under independent purification and assay conditions. All mutants purified successfully with yields comparable to those obtained for the wild-type protein (Table S1).

### Activating mutants cluster at the REP:RING1 and RING0:RING2 interfaces

We initially quantified the E3 ubiquitin ligase activity of human and rat Parkin mutants, using a simple autoubiquitination assay of recombinant Parkin incubated with the E2 ubiquitin-conjugating enzyme UbcH7, as well as the E1 enzyme and ATP. Whilst activating mutants such as W403A robustly autoubiquitinates, it was difficult to quantify Parkin by densitometry because free polyubiquitin chains also formed in the assay and migrated at the same molecular weight as unmodified Parkin (Figures 2A (human) and S1 (rat)). However, we observed that activated Parkin also ubiquitinates UbcH7, whose SDS-PAGE migration is distinct from polyubiquitin chains. We therefore quantified Parkin activity by quantifying the loss of unmodified UbcH7 on SDS-PAGE, as reported by our group recently (Fava et al 2019). All experiments were performed in duplicate and included wild-type and W403A variants on every gel as internal benchmark controls for low and high activity, respectively.

**Figure 2.**
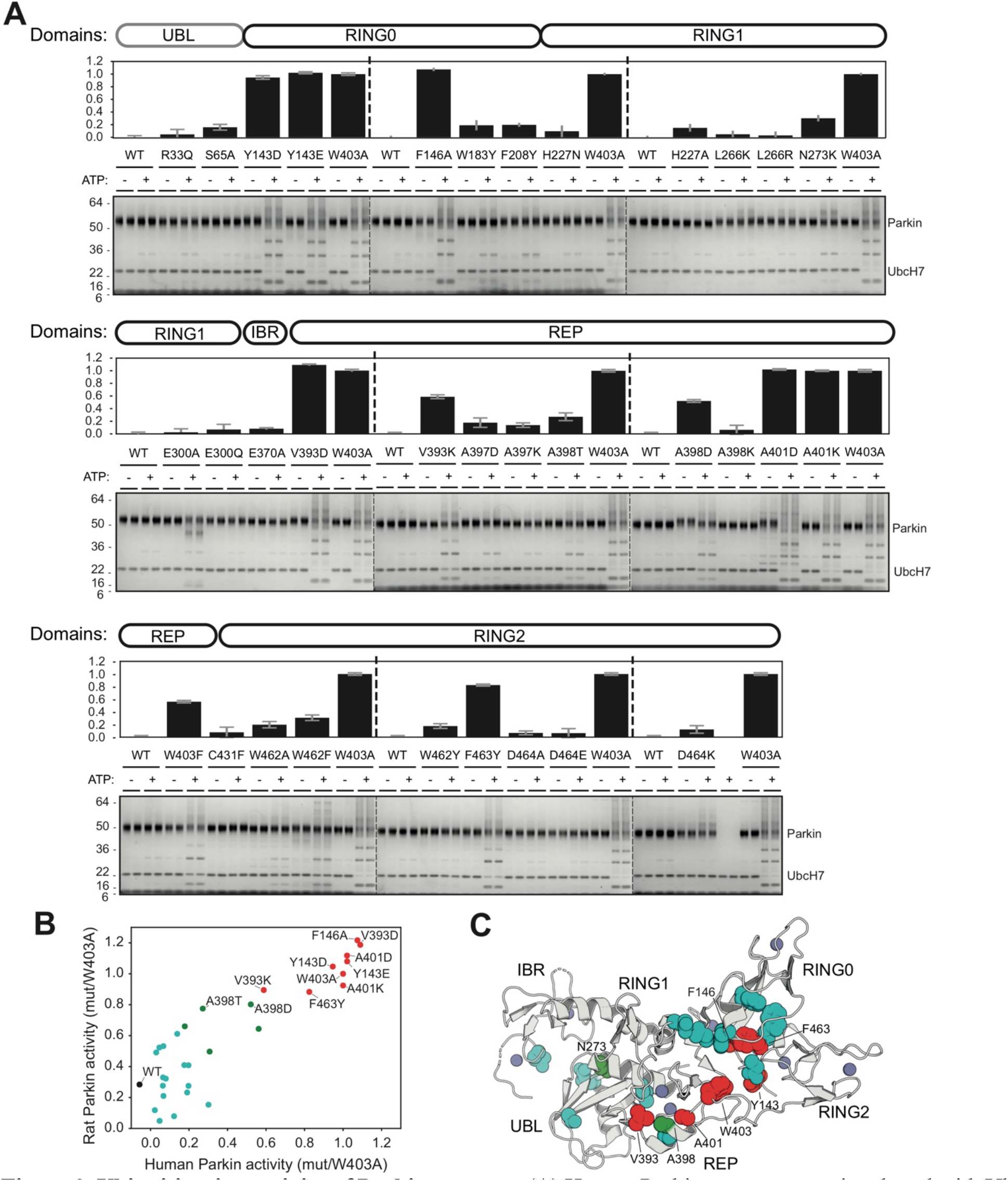
Ubiquitination activity of Parkin mutants. **(A)** Human Parkin mutants were incubated with Ub, E1 and UbcH7 at 37 °C for 1 hour in the presence or absence of ATP. Reactions were resolved on 4-12% Bis-Tris gels followed by Coomassie blue staining (bottom). Activity was determined by measuring the proportion of UbcH7 band intensity lost upon addition of ATP. The bar chart (top) shows the mean depletion from all reactions on each Parkin mutant with error bars indicating the standard deviation. **(B)** Relative activities of human and rat Parkin mutants normalized to the W403A mutant. Dots are coloured by relative activity (red, high; green, medium, cyan, low). **(C)** Structure of autoinhibited rat Parkin (PDB 4ZYN), with spheres indicating the position of mutated amino acids and the colours indicating the relative activity.

There was strong agreement in the activation profile of mutants across human and rat assays (Figure 2B). The data confirms that F146A, W403A, and F463Y activate Parkin in both rat and human in vitro, as observed previously (Riley et al 2013, Trempe et al 2013). We also find that mutants Y143D, Y143E, V393D, A401D, and A401K activate Parkin to levels comparable to W403A in both species. The mutants V393K, A398D, and W403F activated Parkin to a lower extent in both species. Finally, we found that the A398T mutation in human Parkin had a minor effect, but a stronger effect in rat Parkin. Finally, all other mutations had minor or no effects on Parkin ligase activity. Overall, these experiments identify 11 mutations that consistently activate both rat and human Parkin. These mutations are all located at the RING0:RING2 or REP:RING1 interfaces (Figure 2C), suggesting these are most critical for auto-inhibition.

### Thermal stability assays reveal that destabilizing mutations activate Parkin, but only at the RING0:RING2 or REP:RING1 interfaces

Previous studies have suggested that Parkin could be activated through destabilization of its auto-inhibited state (Caulfield et al 2014, Chaugule et al 2011, Riley et al 2013, Spratt et al 2013, Trempe et al 2013, Wauer & Komander 2013). To investigate this hypothesis, we determined the stability of recombinant Parkin mutants using a fluorescence-based thermal shift assay (Figure 3A). WT rat and human Parkin have an average melting temperature (*T_m_*) of 55.8°C and 59.0°C, respectively. Overall, all 11 activating mutations lower the *T_m_* of Parkin by 3-7°C compared to WT. To better understand the relationship between thermal stability and ubiquitin ligase activity, we performed a correlation analysis between the two quantities (Figure 3B). The results for both rat and human Parkin show an inverse correlation between the *T_m_* and the relative ubiquitination activity (Pearson correlation *r*^2^: 0.63 in human, 0.51 in rat). Furthermore, if we restrict the analysis to mutations located at the RING0:RING2 or REP:RING1 interfaces, the correlation is even more significant (*r*^2^: 0.89 in human, 0.80 in rat), whereas there is no correlation for residues located outside this region (Figure S2). For example, mutations at Trp183, Asn227 and Glu300, which are all interacting with each other at the RING0:RING1 interface (Figure 1B), led to destabilization but no activation. Therefore, we conclude that activation is achieved specifically by reducing interdomain interactions that lead to dissociation of the REP (necessary for E2 binding) and the RING0 domain (necessary for thioester transfer on the catalytic RING2 domain).

**Figure 3.**
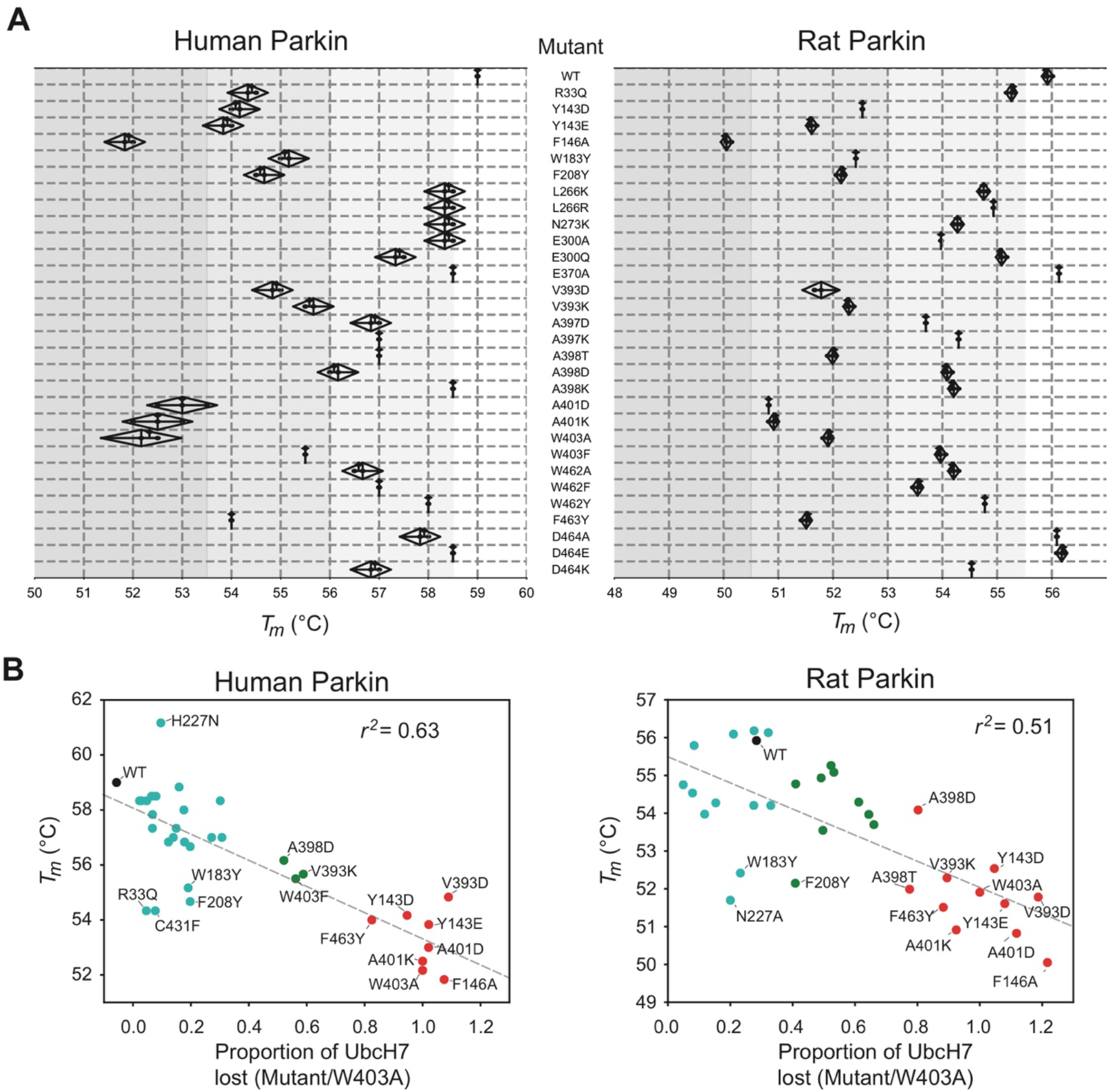
Thermal stability of Parkin mutants and correlation with E3 ligase activity. **(A)** Thermal shift assays for human Parkin (left) and rat Parkin (right). For each mutant, the melting temperature (*T_m_*) was determined by measuring the change in SYPRO-orange fluorescence intensity with temperature. The *T_m_* is the temperature which gives the maximum slope of the temperature against fluorescence. Measurements were made in triplicates, with the mean, maximum and minimum *T_m_* from human and rat Parkin displayed in the above kite plots. **(B)** Scatter plots of the *T_m_* correlated with the E3 ligase activity, measured as loss of UbcH7 and normalized to W403A, for both human (left) and rat (right) Parkin mutants. The dashed line is a linear regression of all the data points, with the correlation coefficient *r^2^* indicated. Dots are coloured by relative activity (red, high; green, medium, cyan, low) as in Figure 2B,C.

### Addition of a negative charge on Tyr143 increases Parkin ubiquitination activity

Our ubiquitination assays revealed that the Y143D and Y143E mutations activate Parkin in vitro (Figures 2 and S1). This result was surprising, given that two groups previously reported that phosphorylation of Tyr143 leads to the inactivation of the ligase activity (Imam et al 2011, Ko et al 2010). Since aspartic acid and glutamic acid lack the aromatic group of a phospho-tyrosine, we therefore sought to test Tyr143 phosphorylation directly. However, we were unable to obtain pTyr143-Parkin; in vitro phosphorylation with recombinant c-Abl led to non-specific, low-level phosphorylation of all solvent-exposed tyrosine residues in Parkin (Figure S3). We therefore incorporated the non-natural phospho-tyrosine mimetic p-carboxymethylphenylalanine at position Tyr143 by using the amber stop codon and modified tRNA and tRNA synthase (Xie et al 2007). Strikingly, the Y143X mutation increased the autoubiquitination activity of Parkin, similarly to the Y143E mutation (Figure 4). Overall this finding indicates that addition of a negative charge at the intersection of RING0, RING1 and the C-terminus results in the release of autoinhibition in Parkin.

**Figure 4.**
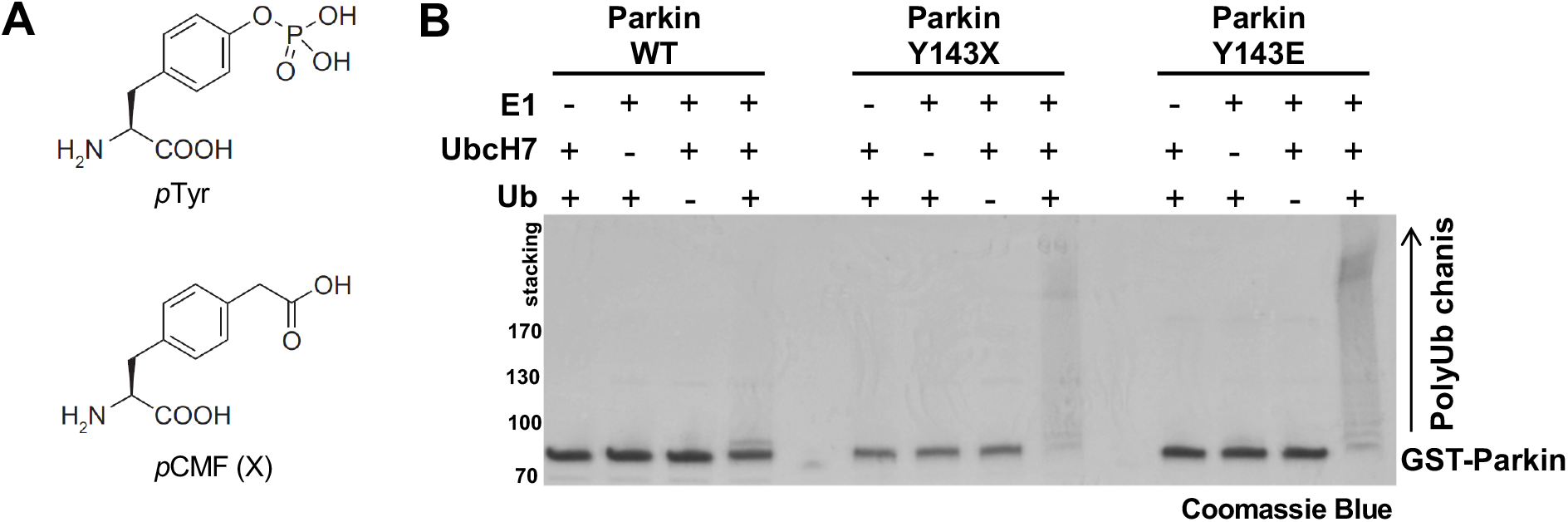
Ubiquitination activity of Parkin Tyr143 phospho-mimetic mutants. **(A)** Chemical structure of phospho-tyrosine and the mimetic artificial amino acid p-carboxymethyl phenylalanine (pCMF). **(B)** 30 min autoubiquitination assay with recombinant GST-fusion rat Parkin wild-type (WT), Y143E and Y143X, where X is pCMF. Reactions were resolved by SDS-PAGE and proteins stained with Coomassie Blue. Note the ubiquitinated ladder (smear) appearing in the presence of ubiquitin, E1 and E2 enzymes, as well as the loss of the unmodified GST-Parkin band, in both Y143E and Y143X mutants, which indicates stronger E3 ubiquitin ligase activity.

### Mutation of Trp403 abrogates binding to the PRK8 monoclonal antibody

To aid in downstream studies. we next determined the best antibody for monitoring levels of Parkin mutants. We therefore performed SDS-PAGE analysis of all recombinant rat Parkin mutants and immunoblotted them against two antibodies. The first is the widely used monoclonal mouse antibody PRK8, directed towards the C-terminus of Parkin (Pawlyk et al 2003). The second one is an in-house polyclonal sheep antibody that recognizes the N-terminus of Parkin (S229D; https://mrcppureagents.dundee.ac.uk/reagents-view-antibodies/589302). The PRK8 antibody robustly detected all mutants except W403A and W403F, implying that Trp403 is a key element of the antibody epitope (Figure S4). In contrast, the N-terminal Parkin antibody recognized all mutants equally well including W403A and W403F (Figure S4).

### Mitophagy assay identifies activating mutants that can rescue S65A Parkin

We next sought to determine whether mutations that increase Parkin ubiquitination activity in vitro also increase mitochondrial depolarization-dependent mitophagy in human cultured cells. Indeed, we had previously observed that the W403A and F146A Parkin mutants recruit faster to mitochondria and increase mitophagy, as measured using the pH-sensitive fluorescent reporter mito-Keima (Katayama et al 2011, Tang et al 2017). We thus introduced 11 activating mutations in GFP-Parkin (human) for transient transfection in U2OS cells stably expressing inducible mt-Keima. Novel mutations that showed the greatest increase in activity in vitro were chosen (Y143E, Y143D, V393D, A401D, A401K, F463Y), as well as V393K, W403A, D464K and E300A for comparison. We also included A398T as it gave contradictory results in rat and human. All mutants expressed to similar levels and showed no accelerated degradation in the presence of the translation inhibitor cycloheximide, suggesting they do not undergo rapid autoubiquitination in cells (Figure S5). The results show that only the V393D and A401D mutants, in addition to W403A, show a significant increase in mitophagy (Figure 5 and S6).

**Figure 5.**
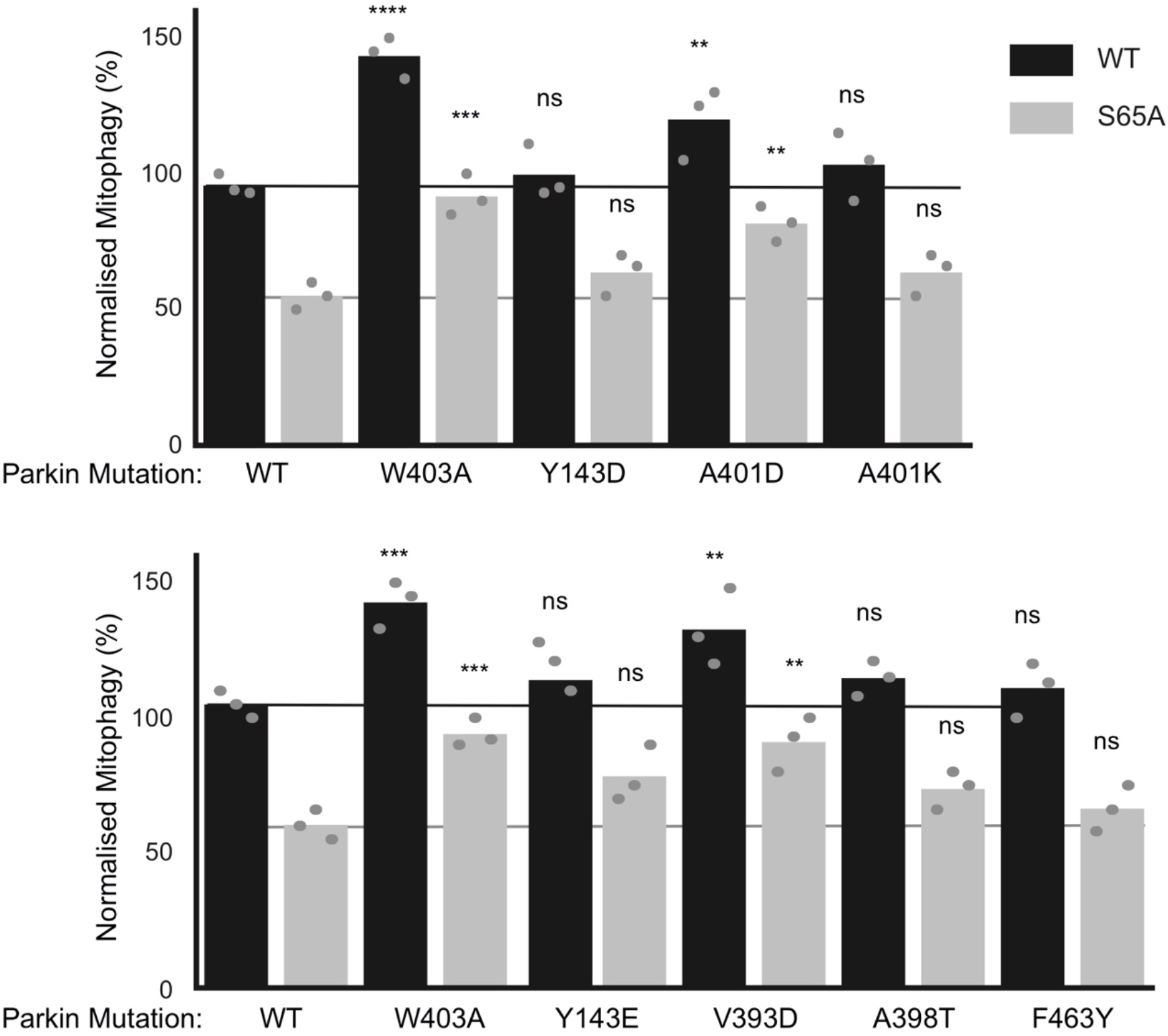
The effect of Parkin mutations on mitophagy. Mitophagy assays were performed in U2OS cells stably expressing mito-Keima and transiently transfected with GFP-tagged human Parkin mutants. Bars indicate the percentage level of mitophagy, normalised to WT, for human Parkin mutants in both WT (black) and S65A (grey) backgrounds. The horizontal lines indicate WT level (dark grey) and S65A level (pale grey line). One-way ANOVA with Dunnett’s post hoc tests (*n* = 3), *P < 0.05; **P < 0.01; ***P < 0.001; ****P<0.0001; ns, not significant.

We next determined which of those mutations would rescue the S65A mutant, which cannot be phosphorylated by PINK1. We had previously observed that the F146A and W403A mutations could rescue the S65A mutation (Tang et al 2017). We therefore combined activating mutations with the S65A and measured normalized mitophagy. We included the mutants that showed a significant elevation of mitophagy (V393D, A401D, W403A), as well as A401K, Y143D/E, and F463Y mutants because they all significantly increased Parkin’s ubiquitination activity in vitro (Figure 2A). The results show that there are only three mutants that can rescue S65A: V393D, A401D, and W403A (Figure 5).

## Discussion

In this study, we engineered 31 mutations across all interdomain interfaces of Parkin including the Ubl:RING1, RING0:RING1, REP:RING1, and RING0:RING2 interactions. This led to the identification of 11 mutations that were capable of activating Parkin in both human and rat species in vitro (Figures 2 and S1). Notably, all mutations were localised near the RING0:RING2 or REP:RING1 interfaces and reduced the thermal stability of Parkin (Figure 3). On the other hand, mutations that destabilised the Ubl, IBR or RING0:RING1 interaction had generally minor or no effect on activation, regardless of their impact on the thermal stability. For instance, mutations in Trp183, Phe208, or Glu300, at the RING0:RING1 interface, destabilise Parkin to various degrees, but do not activate Parkin E3 ligase activity. The mutation R33Q, which has been shown to reduce the thermal stability of the isolated Ubl domain (Safadi et al 2011), also reduce the overall Parkin stability, but yet does not activate the ligase. In other words, reduced stability alone does not predict activation, but all activating mutations destabilise apo Parkin. This explains the strong inverse correlation between the *T_m_* and ligase activity observed for mutations at the REP:RING1 and RING0:RING2 interfaces alone (Figures 3 and S2). This also reinforces the current structural model of Parkin activation, which proposes that both the REP and RING2 must dissociate to allow E2∼Ub binding and thioester transfer of Ub to Cys431 in RING2. Binding of phospho-Ubl to the RING0 domain stabilises this “open” configuration and thus displaces the chemical equilibrium towards the active form. Activating mutations also shift this equilibrium towards the “open” configuration and thus mimic the effect of Ubl phosphorylation.

Our observation that mutations Y143D, Y143E and the phospho-tyrosine mimetic Y143X increase Parkin’s activity (Figure 1, S2 and 4) was not surprising from a structural standpoint, since the sidechain of Tyr143 in RING0 points towards the RING0:RING2 interface in the autoinhibited structure of rat Parkin (Sauvé et al 2015, Trempe et al 2013). The interface notably includes Asp464 as well as the carboxy-terminal residue Val465 in RING2 (Figure 1). Thus, introduction of a negative charge at this position can lead to electrostatic repulsion and dissociation of the interface. It should be noted that the autoinhibited structure of human Parkin (Kumar et al 2015) was determined from a construct with a deletion of the Ubl-RING0 linker that spans a.a. 84-143, and thus Tyr143 is not required to maintain the autoinhibited conformation. However, our results are in stark contrast to those obtained by two groups who reported that Tyr143 phosphorylation by c-Abl inhibits Parkin’s activity in vitro and in vivo (Imam et al 2011, Ko et al 2010). The two groups notably showed that c-Abl can phosphorylate recombinant Parkin at Tyr143, and that the modification can be blocked by Gleevec (STI-571, imatinib), a selective Abl kinase inhibitor. c-Abl was proposed as a target for disease-modifying therapy for PD, with Abl inhibitors like nilotinib displaying neuroprotective activity against 1-methyl-4-phenyl-1,2,3,6-tetrahydropyridine (MPTP)-lesioned mice (Karuppagounder et al 2014). However, we were not able to phosphorylate selectively Tyr143 in vitro using c-Abl, under conditions where PINK1 phosphorylates Ser65 (Figure S3A,B). Recombinant c-Abl was active under these same conditions because it autophosphorylated extensively at several tyrosine residues (Figure S3C). While we cannot exclude that Parkin can be phosphorylated in cells by c-Abl under some conditions, our in vitro results suggest that c-Abl does not selectively phosphorylate Tyr143 in Parkin, and if it does, it would activate Parkin rather than inhibit it. In any case, nilotinib worsened motor symptoms in moderately advanced PD patients and did not modify biomarker outcomes such as dopamine metabolites, suggesting that c-Abl may not be a suitable target for PD treatment (Simuni et al 2021).

In previous work, we found that the F146A and W403A mutations accelerated Parkin recruitment to depolarized mitochondria and increased mitophagy flux (Tang et al 2017). In contrast, the hyperactive mutant N273K, which disrupts the Ubl:RING1 interaction, exhibited slightly increased rates of mitochondrial recruitment and substrate ubiquitination but no alteration in mitophagy. These findings suggested that in cells, disruption of the REP:RING1 or RING0:RING2 interfaces but not the release of the Ubl domain were rate-limiting steps for Parkin activation on mitochondria and initiation of mitophagy. We therefore assessed whether any of the mutations that enhance Parkin activity in vitro could accelerate mitophagy in cells (Figure 5). We found that the V393D and A401D Parkin mutants were able to increase mitophagy flux similarly to the W403A mutation. On the other hand, the mutations Y143D, Y143E, A401K and F463Y, which strongly activate Parkin in vitro (Figure 2), did not increase mitophagy (Figure 5). While these mutants are not destabilised in cells (Figure S5), we cannot exclude that these mutations have additional effects on Parkin-mediated mitophagy that counteract any stimulating effect of its E3 ubiquitin ligase activity. We note that mutations of REP residues Val393 or Ala401 to aspartic acid increased mitophagy, whereas mutation of the same residues to lysine did not (Figures 5 and S6). Furthermore, we observed that the mutations A398T and A398D slightly increased Parkin’s activity in vitro, whereas the A398K mutation did not (Figure 2). It therefore seems that introduction of a positively charged residue in the REP dampens activation, perhaps as a result of the REP making additional, yet unknown interactions in the thioester transfer complex with a charged E2∼Ub conjugate. The structure determination of that complex will help better understand how these mutations activate Parkin.

Autosomal recessive mutations in Parkin represent the most frequent genetic cause of early-onset PD (Klein & Westenberger 2012). The structure-function analysis of Parkin E3 ligase activity to date has been exemplar in understanding the mechanism of how PD mutations disrupt Parkin activation. We have notably reported the utility of the mitophagy assay in assessing 51 naturally occurring variants of Parkin and found that pathogenic variants exhibit severe mitophagy defects in cells, whereas clinically benign variants do not significantly impact mitophagy (Yi et al 2019). Pathogenic variants could be attributed to disruption of different features of Parkin, namely Ubl folding, PINK1 activation mechanism, catalytic activity, and zinc coordination. The two synthetic mutations F146A and W403A, as well as the naturally occurring variant V224A to a lesser extent, were able to rescue in *cis* defects in Ubl folding and most mutations in the PINK1 activation pathway, but not variants that disrupt catalytic activity or unfold the zinc-finger domains. For example, the Ubl folding mutant R42P, as well as the K161N and K211N mutations that prevent activatory pUbl binding to the RING0 domain, are rescued by the F146A and W403A mutations. This is consistent with the same activating mutations being able to rescue deletion of the Ubl or the S65A mutations (Tang et al 2017). Here, we find that the V393D and A401D mutations, in addition to W403A, can also rescue the S65A mutation in *cis* (Figure 5B). On the other hand, the Y143D, Y143E, A398T and F463Y mutations were not able to rescue the S65A mutation. The A398T mutation, which is a naturally occurring variant, did not strongly activate Parkin in vitro (Figure 2). Likewise, the F463Y mutation had a more modest activating effect than the V393D and A401D mutations. Thus, it seems like the ability to rescue the S65A mutant in mitophagy correlates with in vitro E3 ligase activity, with the notable exception of the Tyr143 mutants. In future work, it will be interesting to test the ability of V393D and A401D to rescue pathogenic Parkin mutants in a manner similar to the W403A and F146A mutations.

The Val393, Ala401, and Trp403 residues are all located within the REP domain and play a critical role in mediating the REP:RING1 autoinhibitory interaction. Our findings add to accumulating data suggesting that destabilization of the REP:RING1 interface can circumvent specific functional and pathogenic mutants of Parkin with therapeutic implications. The development of small molecule Parkin enzyme activators has remained extremely challenging in the field to date. Biogen has recently identified a number of compounds that act as positive allosteric modulators of Parkin (Shlevkov et al 2022). These compounds sensitize Parkin to the activating effect of pUb or the W403A mutation but fail to enhance Parkin recruitment or mitophagy. The elaboration of further REP domain activating mutations and their ability to rescue Parkin function in cells, adds further impetus to a review on how future screens of Parkin activators should be performed. Screening should be designed to specifically search for small molecules that mimic REP:RING1 destabilisation both in terms of mechanism of action and amplitude of effect of REP mutants. Were a therapeutic to be developed that mimics REP:RING1 destabilisation, not all Parkin mutants can be rescued including mutations that disrupt catalytic activity or zinc coordination indicating the importance of stratification of Parkin mutant carriers that could benefit from this strategy.

## Materials and Methods

### Reagents

All mutagenesis was carried out using the QuikChange site-directed mutagenesis method (Stratagene) with KOD polymerase (Novagen). cDNA constructs for human Parkin mutant expression were amplified in Escherichia coli DH5*α* and purified using a NucleoBond Xtra Midi kit (#740420.50; Macherey-Nagel). All DNA constructs were verified by DNA sequencing, which was performed by The Sequencing Service, School of Life Sciences, University of Dundee, using DYEnamic ET terminator chemistry (Amersham Biosciences) on Applied Biosystems automated DNA sequencers. DNA for bacterial protein expression was transformed into *E. coli* BL21 DE3 RIL (codon plus) cells (Stratagene). cDNA plasmids (Table 1) and recombinant human Parkin proteins generated for this study are available to request through reagents website https://mrcppureagents.dundee.ac.uk/.

### Recombinant Protein Expression

#### Human Parkin

His_6_-SUMO cleaved wild type Parkin was expressed based upon the method (Kazlauskaite et al., 2014b). Briefly, plasmids were transformed in BL21 Codon Plus (DE3)-RIL *E. coli*, overnight cultures were prepared and used to inoculate 12 x 1 L of LB medium containing 50 μg/ml carbenicillin and 0.25 mM ZnCl_2_. These were initially incubated at 37 °C until the cultures reached an OD_600_ of 0.4, the incubator temperature was lowered to 15 °C, and once cultures reached an OD_600_ of 0.8-0.9 expression was induced by the addition of 25 μM IPTG. After overnight incubation (16 h) cells were pelleted by centrifugation (4200 x g, 25 minutes), the media was removed, and the cell pellet was suspended in lysis buffer [50 mM Tris-HCl pH 7.5, 250 mM NaCl, 15 mM imidazole (pH 7.5), and 0.1 mM EDTA with 1 μM AEBSF and 10 μg/ml leupeptin added]. Cells were burst by sonication and cell debris were pelleted by centrifugation (35000 x g for 30 minutes at 4°C) and the supernatant was incubated with Ni-NTA resin for 1 hour at 5-7°C. Ni-NTA resin was washed 5 times in 7 x the resin volume of lysis buffer, and twice in 7 x the resin volume of cleavage buffer [50 mM Tris pH 8.3, 200 mM NaCl, 10 % glycerol and 1 mM TCEP]. Parkin was cleaved from the resin at 4°C overnight by the addition of a 1:5 mass ratio of His-SENP1 to total protein mass bound to the resin. After cleavage Parkin was concentrated and further purified using size exclusion chromatography on a Superdex S200 column (16/600). Parkin was eluted after 80 to 90 ml, fractions were pooled and concentrated the purity was tested using SDS-PAGE. All the mutants could be concentrated to more than 1 mg/mL. The purity of each mutant was confirmed using SDS-PAGE.

#### Rat Parkin

*Rattus norvegicus* full-length Parkin DNA was codon-optimized for *E.coli* expression and subcloned into pGEX6P-1 (DNAexpress). PCR mutagenesis was used to generate parkin single-point mutants. Parkin expression was done in BL21 (DE3) *E. coli* cells using conditions previously described (Hristova et al 2009, Trempe et al 2013). Cells were grown at 37C to OD ∼1.0, IPTG and ZnSO_4_ were added to 25 μM and cells were left growing overnight at 16°C. Cells were pelleted by centrifugation (3000 x g, 20 minutes), the media was removed, and the cell pellet was suspended in lysis buffer (50 mM Tris/HCl, 120 mM NaCl, 2.5 mM DTT, pH 8.0) with EDTA-free protease inhibitors cocktail (Roche). All proteins were purified using Glutathione-Sepharose 4B (Cytiva) and eluted with 20 mM reduced glutathione in 50 mM Tris/HCl, 120 mM NaCl, 2.5 mM DTT, pH 8.0. Eluted proteins were cleaved overnight with 3C protease followed by gel filtration on Superdex 200 16/60 (GE Healthcare) in 20 mM Tris-HCl, 120 mM NaCl, 2 mM DTT, pH 8.0. All the mutants could be concentrated to more than 1 mg/mL. The purity of each mutant was confirmed using SDS-PAGE.

### Purification of Parkin Y143X mutant

The codon for Tyr143 was mutated to the TAG amber in the pGEX-6p1 rat Parkin plasmid and co-transformed with the pEvol-pCmF plasmid obtained from Peter G. Schultz, as described (Xie et al 2007). Cells were grown overnight in LB medium (20 mL) with chloramphenicol (50 µg/mL) / ampicillin (50 µg/mL) from a single colony. This stock was used to inoculate 500 mL of M9 minimal media containing D-glucose (0.4% w/v), NH_4_Cl (0.1% w/v), MgSO_4_ (4 mM), CaCl2 (0.1 mM), thiamine/vitamin B1 (0.0002% w/v), FeSO4 (4 μM), ZnCl_2_ (4 μM) with chloramphenicol (50 µg/mL) / ampicillin (50 µg/mL) and grown at 37°C until OD600 ∼ 0.5 is reached. Arabinose (0.2%) and pCmF (1 mM) were added and then the culture was grown for another hour. Temperature was reduced to 16°C and equilibrated for 30 minutes. IPTG (25 µM) and ZnCl_2_ (25 µM) was added to induce expression for 18 hours (overnight) at 16 °C. The GST-Parkin-Y143X protein was then purified as described above for rat Parkin.

### Purification of c-Abl and TcPINK1 kinases

Plasmids for human kinase c-Abl (pGEX-cAbl a.a. 83-531) and the YopH phosphatase and was obtained from the lab of John Kuriyan, as described (Seeliger et al 2005). Both plasmids were transformed into BL21-DE3 cells in LB medium with streptomycin (50 µg/mL) / Ampicillin (100 µg/mL). Cells were grown at 37°C until OD600 of 1.2 was reached and cooled for 1 hour at 18° C with shaking. Expression was induced with 0.2 mM IPTG for 16 hours (overnight) at 18°C. Cells were harvested by centrifugation at 7000g at 4°C for 10 minutes and resuspended in 25 mL of cold TBS buffer (50 mM Tris-HCl, 500 mM NaCl, pH 8) with 5% glycerol and EDTA-free protease inhibitors cocktail (Roche). Cells were lysed by sonication and purified using Glutathione-Sepharose 4B (Cytiva) and eluted with 20 mM reduced glutathione in TBS. Eluted protein were resolved by gel filtration on Superdex 200 16/60 (Cytiva) in TBS pH 7.5. GST-TcPINK1 (a.a. 121-570) was purified as described previously (Rasool et al 2018), using a method identical to that described for rat Parkin above, with the exception that no ZnCl_2_ was added.

### Parkin Ubiquitylation Assay

#### Human Parkin

In vitro ubiquitylation assays were performed using recombinant proteins purified from *E. coli* unless stated otherwise. 1 μM of wild type or mutant Parkin was incubated with the ubiquitin master mix [50 mM Tris-HCl, 10 mM MgCl_2_, 2 mM ATP, 0.12 μM His-Ub E1 expressed in Sf21 insect cells, 1 μM human UbE2L3, 50 μM Flag-ubiquitin, and 0.5 μM substrate] to a final volume of 30 μL and the reaction was incubated at 37 °C for 30 min in a thermo shaker at 1000 rpm. Reactions were terminated by the addition of 4 x LDS loading buffer. 5 μl of the final reaction volumes were resolved using SDS-PAGE on a 4-12 % Bis-tris gels in MOPS buffer and stained by incubating with a Coomassie stain (Instant Blue) overnight at room temperature. The stain was washed off using warm water until the gel background had cleared, approximately 3 washes. Gels were imaged using the Licor Odyssey Clx and band intensities were measured using Image Studio Lite. The proportion of UbcH7 depleted was measured by dividing the band intensity of the reaction in the presence of ATP by the band intensity in the absence of ATP and taking this value away from 1. The assay was repeated twice, with the mean UbcH7 levels estimated by densitometry (Figure 2).

#### Rat Parkin

Ubiquitination assays were performed for 2 hours at 37°C. Untagged Parkin (WT and mutant) at 4 μM was incubated with 200 nM E1, 2 μM UbcH7, 100 μM ubiquitin, 0.5 mM TCEP and 10 mM ATP in the presence of 50 mM Tris–HCl pH 7.5 and 5 mM MgCl2. Reactions were stopped with addition of SDS–PAGE sample buffer, resolved on a 12.5% acrylamide Tris/tricine gel containing 0.5% trichloroethanol, and imaged by fluorescence on Gel Doc XR+ Imaging System, (Biorad), as described (Ladner-Keay et al 2018). The assay was repeated twice, with the mean UbcH7 levels estimated by densitometry (Figure S1). For the assay with Parkin-Y143X and Y143E (Figure 4), 1 µM GST-parkin was incubated at 37°C in 50 mM Tris/HCl, 50 mM NaCl, 0.5 mM DTT, pH 7.5, 2 mM ATP, 10 mM MgCl_2_, 40 nM E1, and 2 µM UbcH7, for 60 min. Products were resolved by Tris-glycine SDS-PAGE and the gels stained with Coomassie Blue.

### Thermal Shift Assays

#### Human Parkin

Master mixes containing 1 x SYPRO Orange and 0.06 mg/ml Parkin in the final protein buffer (50 mM Tris pH 8.3, 200 mM NaCl, 10% glycerol and 0.5 mM TCEP) were made up for each Parkin mutant. 50 μl of the master mix was aliquoted into 3 wells of a 96 well PCR plate. The assay was performed using a qPCR machine, for the assay a temperature gradient from 20 to 90 °C was applied in 0.5 °C steps with samples incubated at each step for 1 minute. The intensity of SYPRO Orange emission (570 nm) was recorded at each temperature step. The Tm was determined by taking the first derivative of the SYPRO Orange emission curve and determining the temperature this value reached its maximum.

#### Rat Parkin

Stability of recombinant Parkin mutants was determined by fluorescence-based thermal shift assay. Parkin (wt and mutants) was mixed with SYPRO orange protein gel stain (Molecular Probe – Life Technologies) to final concentration 0.5 mg/ml protein and 6x dye and heated from 10 to 70 °C with fluorescence reading every 0.1 °C (4 to 5 sec) in a QuantStudio^TM^ 7 Flex Real-Time PCR System (Life Technologies). Protein thermal melting curve were generated using the Protein Thermal Shift ^TM^ software. Four replicates were performed. The Tm was determined by taking the first derivative of the SYPRO Orange emission curve and determining the temperature this value reached its maximum.

### Phosphorylation assay and mass spectrometry

Rat Parkin WT (4 μM) was incubated with 1 μM GST-Abl or GST-TcPINK1 for 30 min at RT with 2 mM ATP in 50 mM Tris–HCl pH 7.5 and 5 mM MgCl_2_. Reaction samples were diluted in denaturing buffer (3 M urea, 25 mM TEAB pH 8.5, 0.5 mM EDTA) and reduced using 2 mM TCEP for 10 min at 37 °C, followed by alkylation with 50 mM chloroacetamide for 30 min at room temperature in the dark. Chloroacetamide was used to avoid iodoacetamide-induced artefacts that mimic ubiquitination (Nielsen et al 2008). Samples were diluted with 50 mM TEAB pH 8.5 to 1 M urea and digested with 0.5 µg trypsin (Sigma) for 3 h at 37 °C. Digested peptides were purified using C18 Spin Columns (ThermoFisher) and resuspended in 0.1% formic acid. Peptides (0.5 µg) were captured and eluted from an Acclaim PepMap100 C18 column with a 2 h gradient of acetonitrile in 0.1% formic acid at 200 nl/min. The eluted peptides were analysed with an Impact II Q-TOF spectrometer equipped with a Captive Spray nanoelectrospray source (Bruker). Data were acquired using data-dependent automatic tandem mass spectrometry (auto-MS/MS) and analysed with MaxQuant using a standard search procedure against a custom-made FASTA file including rat Parkin, GST-Abl, and GST-TcPINK1. Methionine oxidation and Ser/Thr/Tyr phosphorylation were included as variable modifications. Cysteine carbamylation was included as fixed modification. For intact mass spectrometry, reaction samples were diluted in 1% formic acid and 1 μg was injected on a Waters C4 BEH 1.0/10-mm column and washed 5 min with 4% acetonitrile, followed by a 10-min 4 to 90% gradient of acetonitrile in 0.1% formic acid, with a flow rate of 40 μL/min. The eluate was analyzed on a Bruker Impact II Q-TOF mass spectrometer equipped with an Apollo II ion funnel electrospray ionization source. Data were acquired in positive-ion profile mode, with a capillary voltage of 4,500 V and dry nitrogen heated at 200 °C. Spectra were analyzed using the software DataAnalysis (Bruker). The multiply charged ion species were deconvoluted at 5,000 resolution using the maximum entropy method.

### Immunoblots

Parkin WT and mutants were immunoblotted with two Parkin antibodies. One ug protein was loaded on 10% acrylamide Tris-glycine gel followed by transfer to nitrocellulose membranes. Membranes were blocked followed by overnight incubation at 4°C with primary antibodies: C-terminal Parkin antibody (PRK8; Cell Signaling, 1:40,000 dilution) and N-terminal Parkin antibody (https://mrcppureagents.dundee.ac.uk/reagents-view-antibodies/589302; sheep poly-clonal antibody S229D, 1:240 dilution). HRP-conjugated secondary antibody (HRP-link anti-mouse IgG, Cell Signaling, 1:10 000 dilution for C-terminal Parkin Ab and HRP-link anti-sheep IgG, Sigma, 1:10 000 dilution for N-terminal Parkin Ab) was incubated one hour at RT and protein visualized using Clarity^TM^ Western ECL Substrate (Bio-Rad).

### Mitophagy assay

Mitophagy was examined using a FACS-based analysis of mitochondrially targeted mKeima as previously described (Tang et al 2017, Yi et al 2019). Briefly, U2OS cells stably expressing an ecdysone-inducible mt-Keima were induced with 10 μM ponasterone A, transfected with GFP-Parkin (Wild-type and/or indicated mutants) for 24 h and treated with 20 μM CCCP for 4 h. For FACS analysis, cells were trypsinized, washed and resuspended in PBS prior to their analysis on a LSR Fortessa (BD Bioscience) equipped with 405 and 561 nm lasers and 610/20 filters. Measurement of lysosomal mitochondrially targeted mKeima was made using a dual excitation ratiometric pH measurement where pH 7 was detected through the excitation at 405 nm and pH 4 at 561 nm. For each sample, 50,000 events were collected, and single, GFP-Parkin-positive cells were subsequently gated for mt-Keima. Data were analysed using FlowJo v10.1 (Tree Star). For statistical analysis, the data represent the average percentage of mitophagy from three independent experiments, and *P* values were determined by one-way ANOVA with Dunnett’s post-hoc tests were performed. * P<0.05, **P<0.01, ***P<0.001.

### Cycloheximide pulse-chase assay

Cycloheximide Pulse-chase assay and Western Blotting Analysis

24 h after transfection, 5 × 10^5^ cells were treated with 100 μg/ml CHX for the indicated amounts of time. Cells were harvested and then lysed in 150 μl of lysis buffer (50 mm Tris, pH 7, 8% glycerol (v/v), 0.016% SDS (w/v), 0.125% β-mercaptoethanol (v/v), 0.125% bromophenol blue (w/v), 1 mm PMSF, and 1 μg/ml leupeptin). The samples were sonicated and then resolved by SDS-PAGE on 10% gels along with Precision Plus All Blue protein prestained standards (Bio-Rad). After SDS-PAGE, proteins were transferred onto nitrocellulose membranes (LI-COR Biosciences). The membranes were blocked with 2.5% fish skin gelatin (Truin Science) in 1× PBS with 0.1% Triton X-100), probed with primary and secondary antibodies, and imaged with an Odyssey® infrared imaging system using the manufacturer’s recommended procedures (LI-COR).

## Acknowledgements

This work was supported primarily by a research grant from the Michael J. Fox Foundation (#14681 to JFT, MMKM, EAF, TD, WS), as well as a Wellcome Trust Senior Research Fellowship in Clinical Science (#210753/Z/18/Z to MMKM), EMBO YIP Award (MMKM), Canada Foundation for Innovation (#229792 to JFT), and Canada Research Chair grants (#950-229792 to JFT and #950-232176 to EAF). MAE is a CIHR-Banting Fellow and supported by Parkinson Canada. WS is supported by NIH R01 NS085070. We are grateful to the sequencing service (School of Life Sciences, University of Dundee); Axel Knebel for expression and generation of ubiquitin reagents (MRC PPU); and MRC PPU Reagents and Services antibody teams (co-ordinated by James Hastie).

## Author Contributions

MUS, NC, MAE, ZWZ, TD performed experiments and analysed data. WS, EAF, MMKM and JFT supervised experiments and analysed results. JFT and MMKM wrote the paper with contribution from all the authors. EAF, MMKM, and JFT conceived and supervised the project.

---

**Figure S1.**
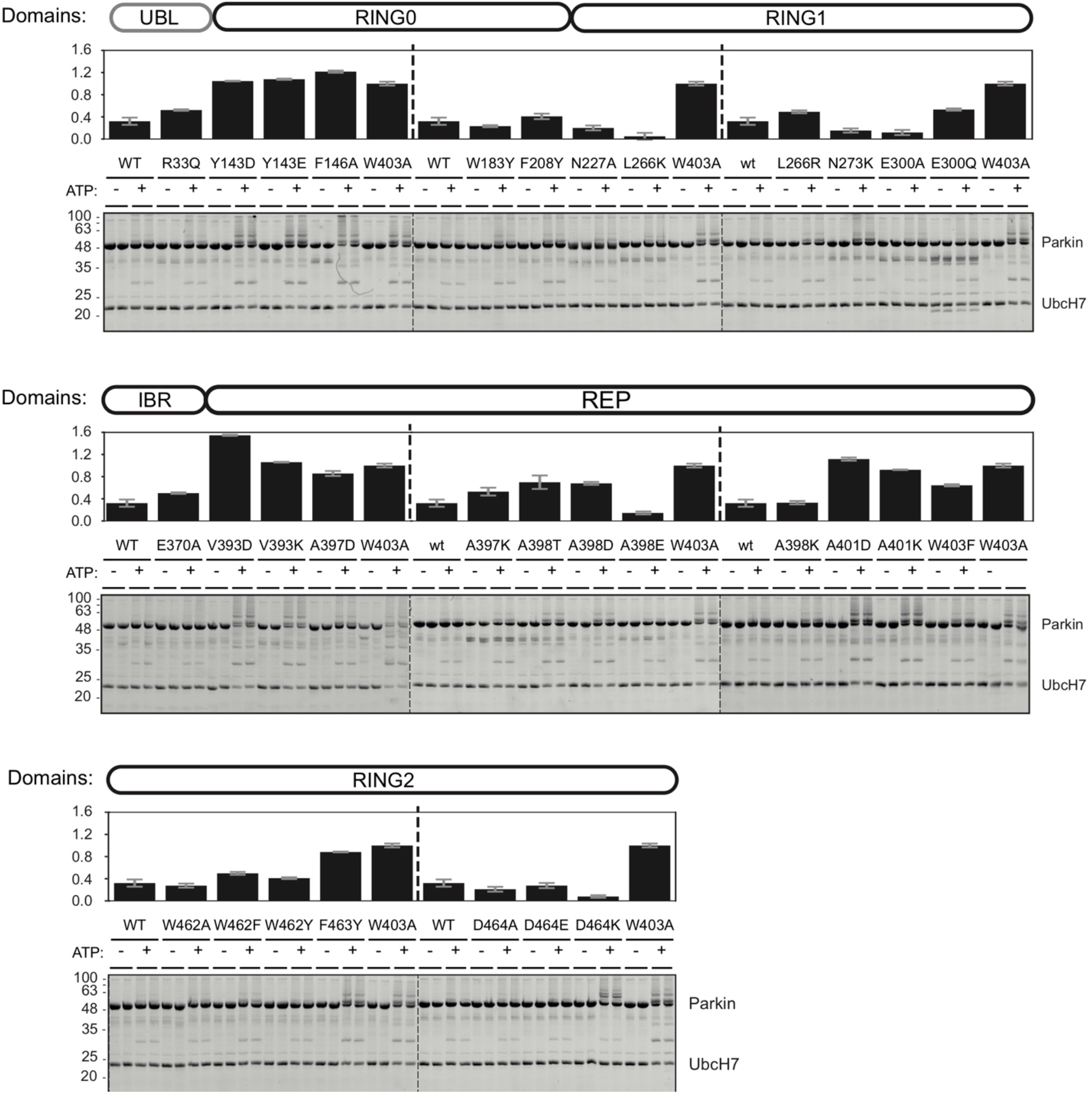
Ubiquitination activity of rat Parkin mutants. Recombinant rat Parkin mutants were incubated with Ub, E1 and UbcH7 at 37 °C for 1 hour in the presence or absence of ATP. Reactions were resolved on Tris-Tricine gels and imaged using TCE and UV fluorescence (bottom). Activity was determined by measuring the proportion of UbcH7 band intensity lost upon addition of ATP. The bar chart (top) shows the mean depletion from all reactions on each Parkin mutant with error bars indicating the standard deviation (*n*=2).

**Figure S2.**
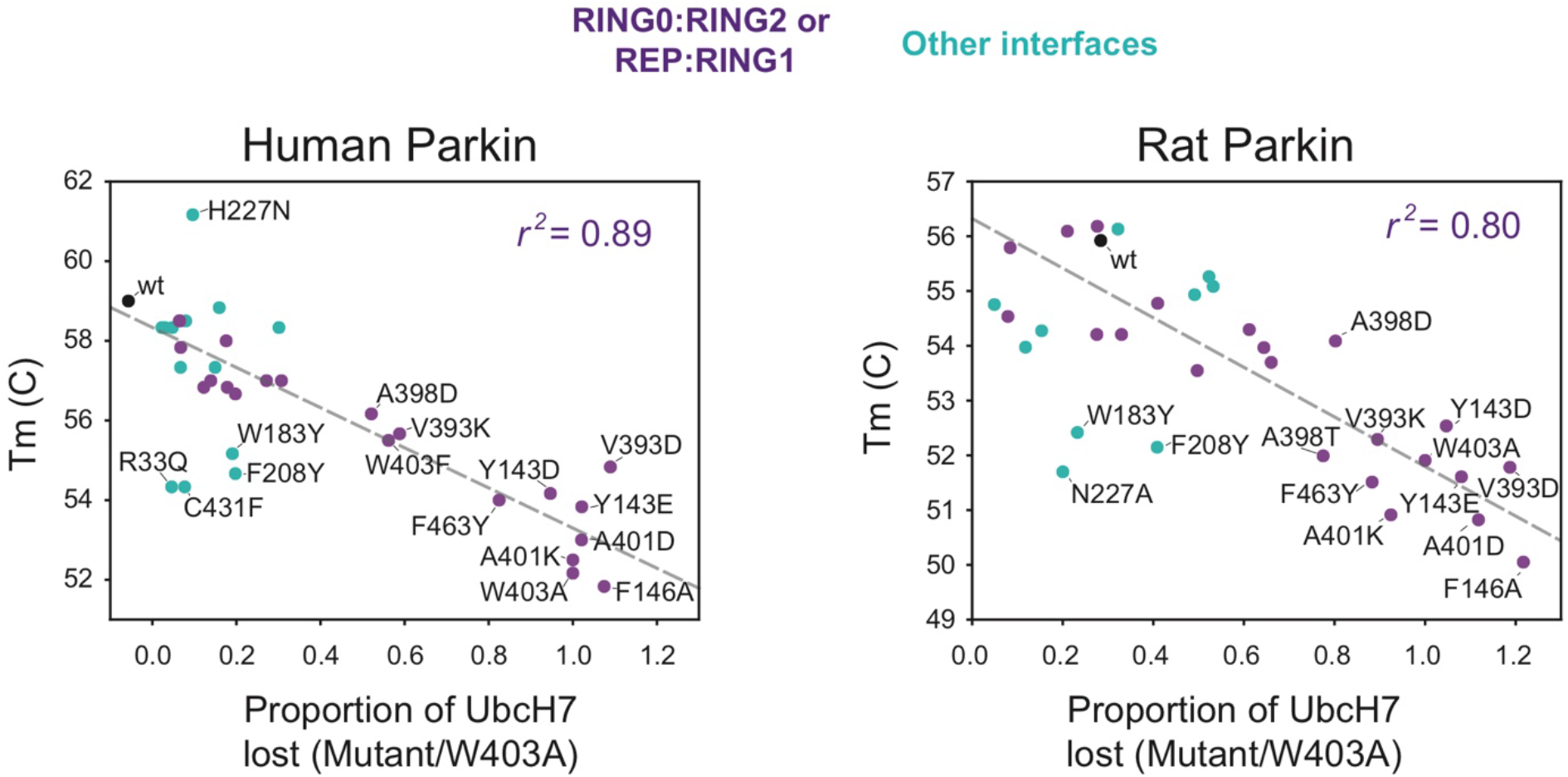
Thermal stability of Parkin mutants and correlation with E3 ligase activity. Scatter plots of the *T_m_* correlated with the E3 ligase activity, measured as loss of UbcH7 and normalized to W403A, for both human (left) and rat (right) Parkin mutants (same data as in Figure 3). Dots are coloured by the location of the mutations, with violet dots indicating residues at the RING0:RING2 or REP:RING1 interfaces, and cyan dots all the other (WT in black). The dashed line is a linear regression of only the violet and black dots, with the correlation coefficient *r^2^* indicated.

**Figure S3.**
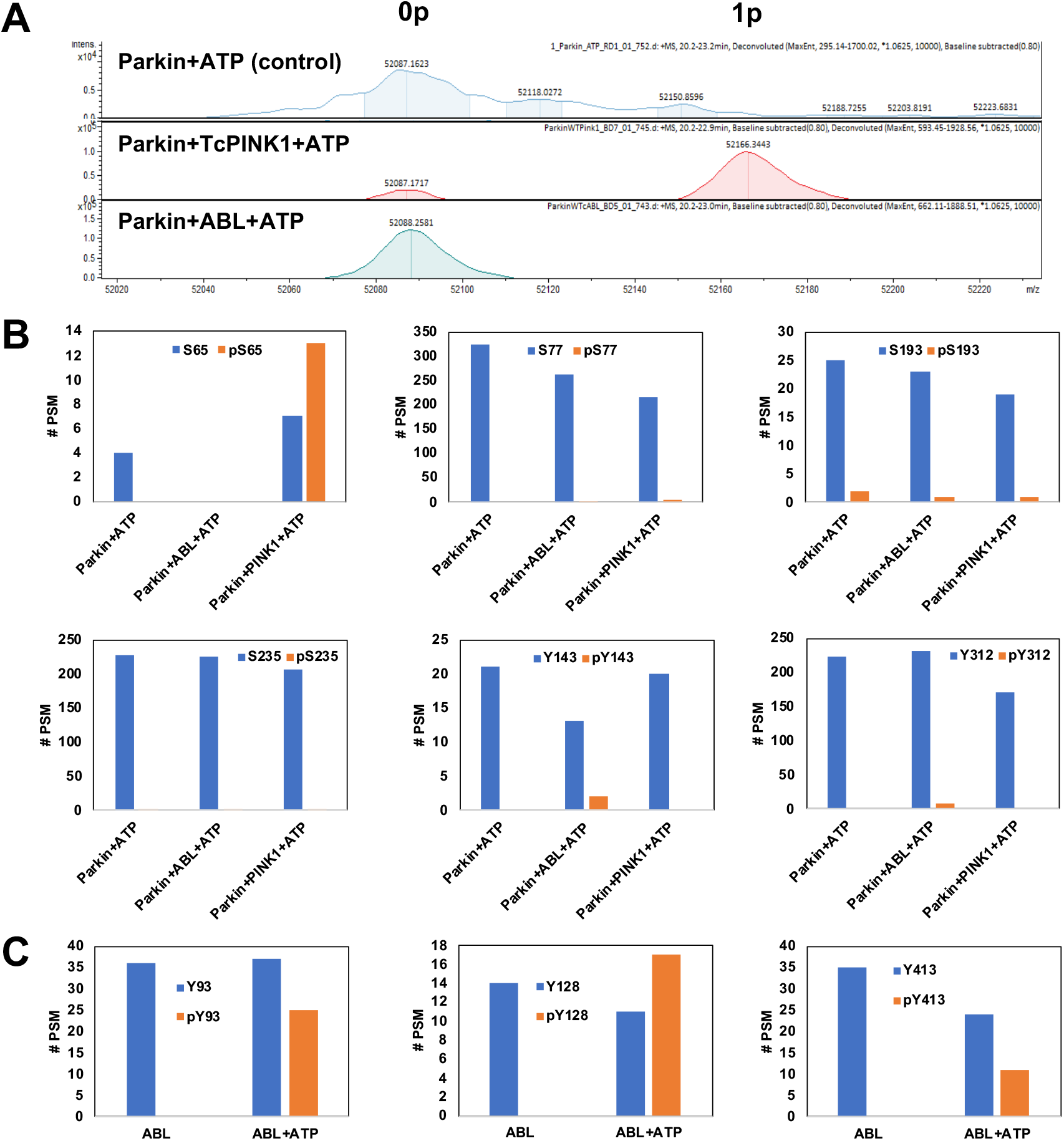
Phosphorylation of Parkin by c-Abl and PINK1. **(A)** Deconvoluted ESI intact mass spectra of recombinant rat Parkin (4 μM, predicted mass 52090.1 Da) incubated 30 min alone, or with 1 μM human c-Abl (a.a. 83-531) or 1 μM PINK1 from *Tribolium castaneum* (TcPINK1 a.a. 121-570), in the presence of 2 mM ATP. The 79.9 Da shift observed upon addition of TcPINK1 indicate addition of a single phosphate (1p). No phosphorylated species appear in the presence of Abl. **(B)** Peptide spectral match # (PSM) for tryptic digestion of the reactions described in A. Each graph corresponds to a peptide for which a phosphorylation site could be detected (blue bar, non-phospho PSM; orange bar, phospho PSM). The data shows that TcPINK1 phosphorylates extensively and specifically Ser65, whereas Abl phosphorylates only a small fraction of Tyr143 and Tyr312. **(C)** Peptide spectral match # (PSM) for tryptic digestion of recombinant Abl incubated in the absence of presence of 2 mM ATP. The data shows substantial autophosphorylation of Abl at Tyr93, Tyr128, and Tyr413, indicating that the enzyme is active.

**Figure S4.**
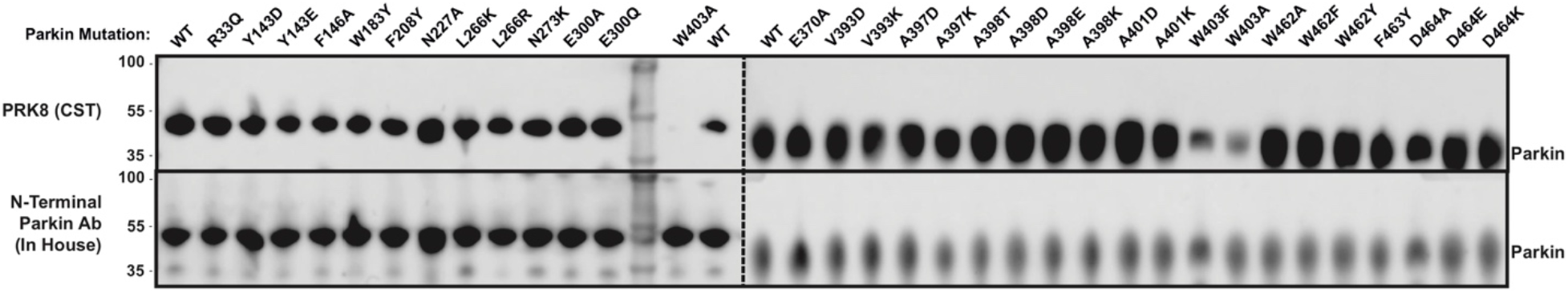
Antibody reactivity of Parkin mutants. Western blots of different Parkin antibodies where the antigen is either the C-terminal region of Parkin including W403 (Cell signalling technology, PRK8, mouse monoclonal), or the N-terminal of Parkin (in house antibody, rabbit polyclonal). Antibodies were tested against different rat Parkin mutations used in this paper.The data shows loss of reactivity of the PRK8 antibody against the W403A and W403F mutants.

**Figure S5.**
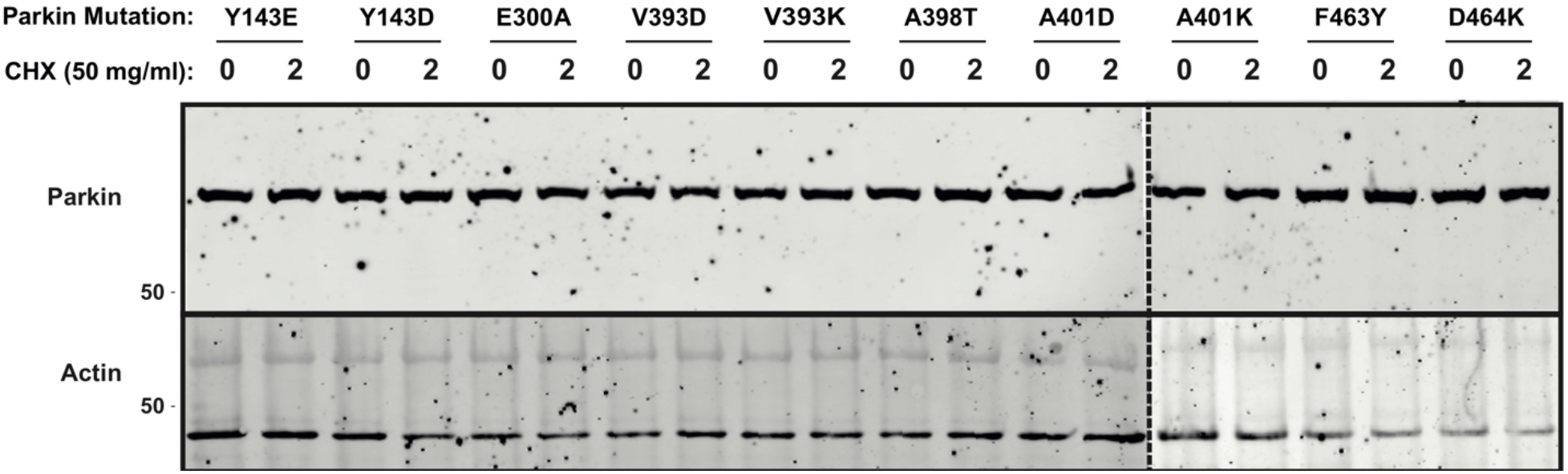
Expression levels and stability of Parkin activation mutants in human cells. Western blots of the different human GFP-Parkin mutants transiently transfected into U2OS cells. Cells were lysed before treatment with the translation inhibitor cyclohexamide, or after treatment for 2 hrs. Loading was checked using actin (bottom blot), with none of the Parkin mutations showing an observable increase in turnover rate compared to wild type (top blot).

**Figure S6.**
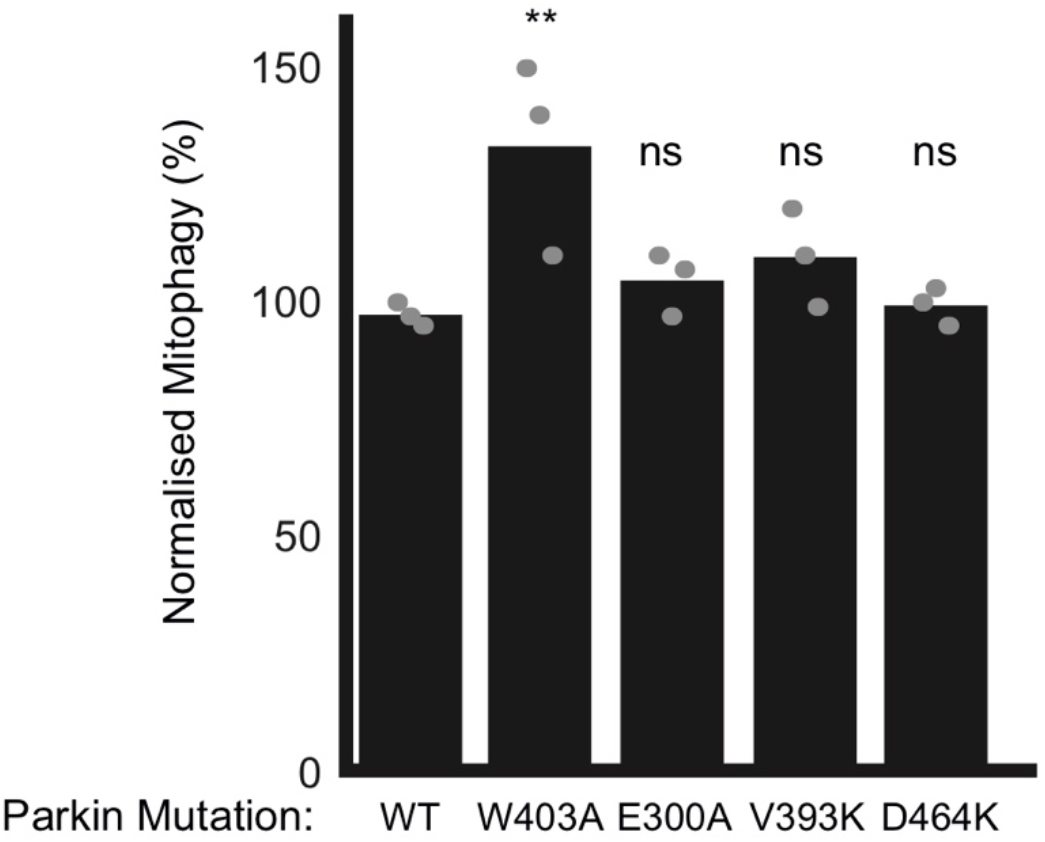
The effect of Parkin mutations on mitophagy. Mitophagy assays were performed in U2OS cells stably expressing mito-Keima and transiently transfected with GFP-tagged human Parkin mutants. Bars indicate the percentage level of mitophagy, normalised to WT, for human Parkin mutants. One-way ANOVA with Dunnett’s post hoc tests (*n* = 3), *P < 0.05; **P < 0.01; ns, not significant.

**Table S1.**
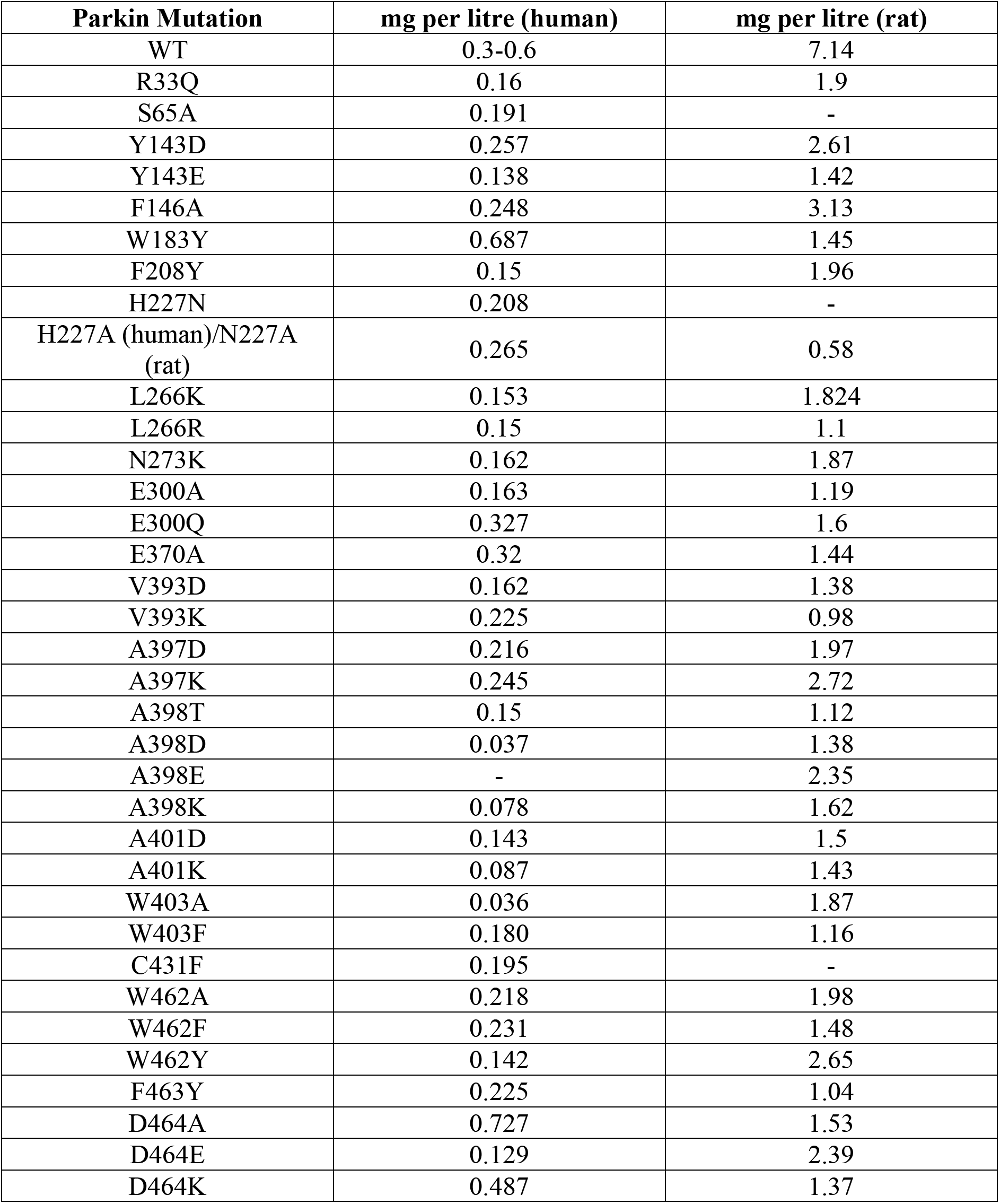
Yields of recombinant Parkin mutants, expressed in E. coli

